# Benchmarking Genetic Interaction Scoring Methods for Identifying Synthetic Lethality from Combinatorial CRISPR Screens

**DOI:** 10.1101/2025.03.31.645224

**Authors:** Hamda Ajmal, Sutanu Nandi, Narod Kebabci, Colm J. Ryan

**Author notes:** To whom correspondence should be addressed. Correspondence may also be addressed to.

## Abstract

Synthetic lethality (SL) is an extreme form of negative genetic interaction, where simultaneous disruption of two non-essential genes causes cell death. SL can be exploited to develop cancer therapies that target tumour cells with specific mutations, potentially limiting toxicity. Pooled combinatorial CRISPR screens, where two genes are simultaneously perturbed and the resulting impacts on fitness estimated, are now widely used for the identification of SL targets in cancer. Various scoring methods have been developed to infer SL genetic interactions from these screens, but there has been no systematic comparison of these approaches. Here, we performed a comprehensive analysis of 5 scoring methods for SL detection using 5 combinatorial CRISPR datasets. We assessed the performance of each algorithm on each screen dataset using two different benchmarks of paralog synthetic lethality. We find that no single method performs best across all screens but identify two methods that perform well across most datasets.

**GRAPHICAL ABSTRACT:** **Figure 1.**
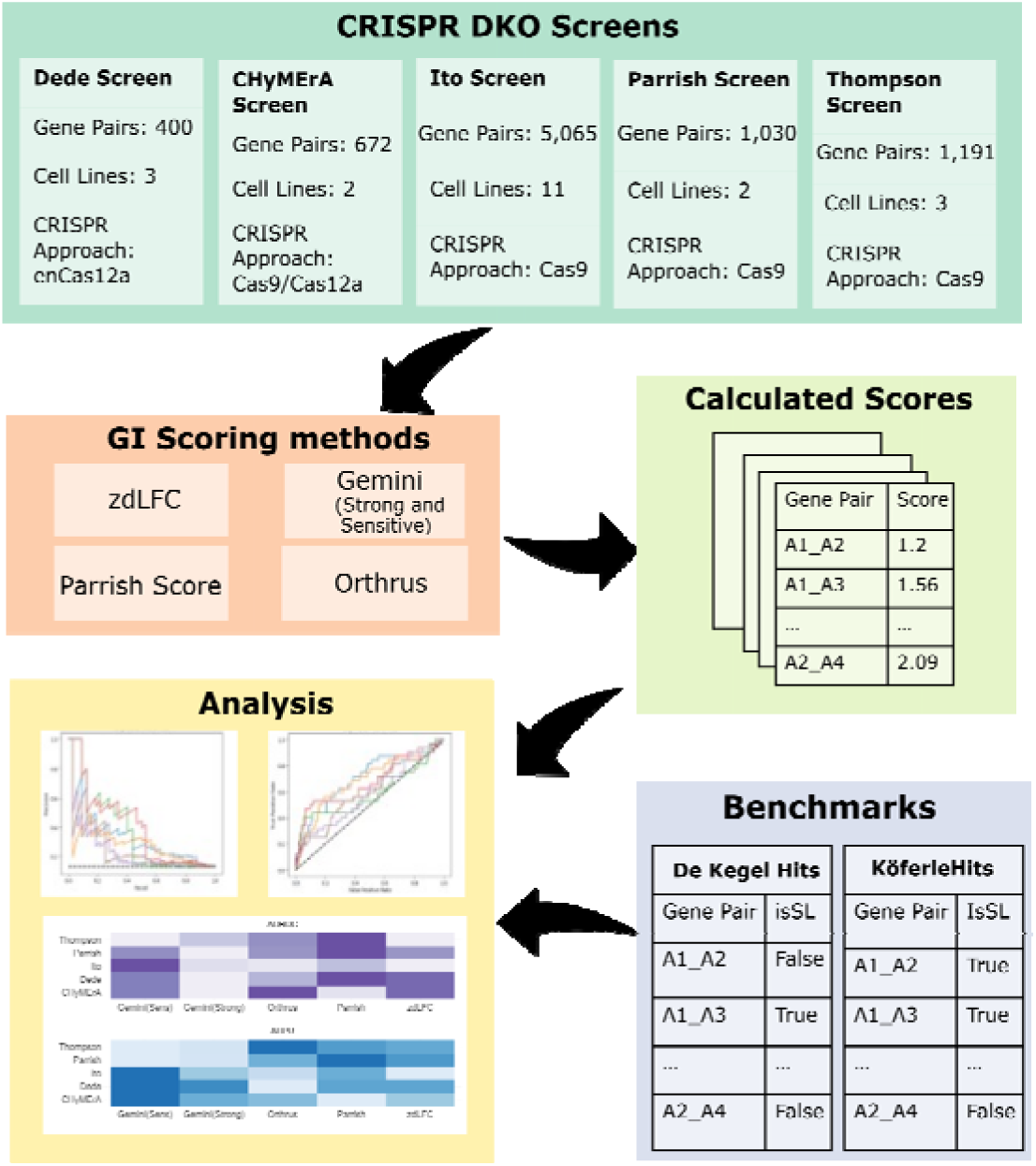
Graphical Abstract. Benchmarking Scoring Methods for Synthetic Lethality Detection from CRISPR screen data. Experimental setup for benchmarking experiments. Five different CRISPR double knockout (DKO) screens are scored for genetic interaction using 5 different scoring methods. The calculated scores are analysed using two different benchmarks (De Kegel Hits and Köferle Hits). Area under the receiver operating characteristic curve (AUROC) and Area under the precision recall curve (AUPR) for each scoring method on each dataset are calculated and compared.

## INTRODUCTION

Synthetic Lethality (SL) is a phenomenon whereby the simultaneous perturbation of a pair of non-essential genes results in cell death (1). Since gene loss, via loss-of-function mutation or deletion, is frequently observed in tumour cells, SL can be leveraged to selectively target these cells, thereby minimizing toxicity to healthy cells. In recent years, extensive research has focused on exploiting SL to develop new cancer therapies (2, 3). A significant milestone in this field was achieved with the approval of the first SL based therapy – using PARP1 inhibitors for treating select breast, ovarian and prostate cancer patients whose tumours have BRCA1 or BRCA2 mutations (2, 4).

CRISPR-Cas (clustered regularly interspaced short palindromic repeats – CRISPR associated protein) screening technology is widely used for genome editing (5) and its advent sparked a renewed interest in finding SL in cancer (6). It uses a guide RNA (gRNA) to identify a target DNA sequence and direct the Cas nuclease to it, where it induces double strand break and causes disruption (5). CRISPR double knockout (CDKO) experiments allow us to examine the impact of simultaneously knocking out pairs of genes to study genetic interactions (GI) between them. They have been widely used to identify synthetic lethal interactions, especially between duplicate genes (paralogs) (1, 7–11). Typically, CDKO libraries contain paired single gRNAs (sgRNAs) knocking out two genes at once. These libraries also include sgRNAs paired with a non-targeting control sgRNA to assess the effects of knocking out each single gene independently. Some experiments may also include two non-targeting paired sgRNAs as a baseline control. Guide RNA (gRNA) abundance is measured through high throughput sequencing at an initial time point and again at later time point(s) following lentiviral transduction.

Analysing and interpreting CDKO screen data remains challenging due to significant variations in guide activity, high replicate variability, and differences in sgRNA library design (12). Variations in sgRNA libraries, such as the number of distinct sgRNA pairs per gene pair, the number of individual or paired non-targeting control sgRNAs, the number of sgRNAs targeting positive control genes and the orientation of targeted gene pairs, make it difficult to apply statistical methods consistently across studies without substantial modifications or adaptations. Several statistical methods (8, 9, 12, 13) have been published in the literature to quantify the magnitude of SL between gene pairs from CDKO screen data. However, most published screens use distinct scoring systems, making comparisons across scoring methods difficult. Currently, there is no consensus on which genetic interaction scoring method yields the best results.

Only a handful of studies have systematically evaluated synthetic lethality scoring methods, and their scope remains limited. Zamanighomi et al. (12), who developed Gemini, evaluated multiple scoring methods but were limited by the lack of a benchmark of true positive and true negative pairs. Moreover, fewer CDKO datasets and scoring systems were available at the time of their publication. Chou et al. (14) compared performance of the dLFC (delta Log Fold Change) method and a number of variants on data from four in4mer screens (15) as well as the (8–10). They did not compare against alternative scoring systems such as Gemini (12) or Orthrus (13), which cannot be directly applied to the in4mer screens. Rather than employing an orthogonal benchmark of synthetic lethality, as we do here, they evaluated the consistency of SL hits identified across screens using the Jaccard coefficient. The Synthetic Lethality Knowledge Base (SLKB) (16) project consolidates data from multiple CDKO experiments scored using various GI scoring methods. However, SLKB lacks comparative insights into the performance of these methods when evaluated against a ground truth. In contrast, our study evaluates the performance of multiple scoring methods and provides a comparative analysis.

In this study, we evaluate multiple GI scoring methods using two benchmarks and rate them based on their performance using Area under the receiver operating characteristic curve (AUROC) and Area under the precision recall curve (AUPR) (Figure 1). We seek to directly address the question of which GI scoring method performs best. We apply 5 scoring methods to 5 CDKO datasets. As benchmarks, we use two different datasets: (1) De Kegel Hits (17), and (2) Köferle Hits (18), detailed in Section ‘Benchmark Datasets’. AUROC and AUPR are computed for each dataset and scoring method combination, and the results are compared across screens.

Overall, our results suggest that the Gemini-Sensitive scoring method and the Parrish score rank consistently higher compared to other methods across all cell lines and both benchmark datasets.

## MATERIAL AND METHODS

### CRISPR DKO studies

In pooled CRISPR screens, the fitness associated with a given construct (gRNA or gRNA pair) is typically inferred from the log-fold change (LFC) in abundance of that construct between early and late time points, with constructs that cause a growth defect having a negative LFC. Typically, a given gene will be targeted with multiple constructs, and an overall gene effect then inferred using the LFC of all these constructs (referred to as ‘single mutant fitness’ or SMF in this paper). In a combinatorial screen a pair of genes will be targeted by constructs containing two gRNAs, and the effect of these paired constructs is used to infer “double mutant fitness” or DMF.

We use datasets from five CDKO studies, detailed in Table 1. Each of these screens uses constructs targeting pairs of genes to study DMF and constructs targeting single genes along with a non-targeting control to study SMF. All screens are pooled combinatorial studies. Read counts are measured at both an early and later time point(s).

**Table 1.** List of CDKO datasets used in this study, along with their details.

| Study Name | CRISPR Approach | Cell Lines | # Gene Pairs | Early Time Point (ETP)/Control | Late Time Point (LTP) |
| --- | --- | --- | --- | --- | --- |
| Dede | enCas12a | A549, HT29, OVCAR8 | 400 | Plasmid pool | 10 population doublings |
| CHyMErA | Cas9 - Cas12a | HAP1, RPE1 | 672 | Day 2 | Day 18(HAP1), Day 24 (RPE1) |
| Ito | Cas9 | Meljuso, GI1, MEL202, PK1, MEWO, HS944T, IPC298, A549, HSC5, HS936T, PATU8988S | 5,065 | Plasmid | Day 21 |
| Parrish | Cas9 | PC9, HeLa | 1,030 | Immediately postinfection and puromycin selection | 12 population doublings |
| Thompson | Cas9 | MEWO, A375 and RPE | 1,191 | Day 7 read counts from wildtype A375 cells (Cas9-negative) transduced with the library under similar screening conditions. | Day 28 |

### Genetic Interaction Scoring Methods

A synthetic lethal relationship is inferred when the observed DMF is significantly worse than the expected DMF, which is typically calculated as the product of the SMF of each single-gene knockout, equivalent to their sum in log space (Mani et al. 2008). We use five different GI scoring methods for our analysis: zdLFC, Gemini-Strong, Gemini-Sensitive, Orthrus and Parrish score (Table 2). These scoring systems primarily differ in how they calculate the expected DMF, and in a variety of other technical details (Table 2).

**Table 2:** List of GI scoring methods analysed in this study, along with their details.

| Scoring Method | Default Filtering | Default pre-processing | Description | Implementation |
| --- | --- | --- | --- | --- |
| zdLFC (8) | - | A pseudo count of 5 is added to all read counts, followed by normalization to an average of 500 reads per guide | Genetic interaction is equal to expected DMF minus the observed DMF. The differences are z-transformed after truncating the top and bottom 2.5% values. $zdLFC \leq -3$ indicates SL-Hit. | The code is provided as Python notebooks, which can be adapted to different datasets |
| GEMINI – Strong (12) | - | Read counts are normalised to the total number of counts. Normalised read counts are adjusted in a way that that median read count across all guides and all genes is set to 0. A pseudo count of 32 is added. | Expected LFC is modelled as a function of sample dependent as well as sample independent individual effects of each guide in a pair and the effect of two guides in combination. Uses the coordinate ascent variational inference (CAVI) approach to infer single and double gene effects. The strong variant identifies interactions where the inferred combination effect significantly exceeds either of the individual effects. Captures genetic interactions with “high synergy”. | Available as an R package with a comprehensive user guide. |
| GEMINI– Sensitive (12) | Removes gene pairs of which an individual gene KO results in more than 50% depletion, compared to gene pairs with the strongest depletion in the screen. |  | The sensitive variant compares the total effect (sum of individual gene effects and the combination effect) with the most lethal individual gene effect. This score can capture “modest synergy”. |  |
| Orthrus (13) | Removes gRNAs with read count < 30 and > 10,000. Gene pairs for which there are too few guides remaining post filtering are also removed. | All counts are log2-scaled (scaling factor 1e6) and normalised to the total number of read counts.<br>A pseudo count of 1 is added post normalisation and scaling. | Assumes an additive linear model for the expected LFC of each gene pair in both orientations (A-B/B-A). Estimates effect size by comparing the expected LFC to the observed LFC for each orientation. Can be configured to ignore orientation when needed. | Available as an R package with a comprehensive user guide. |
| Parrish Score (9) | gRNAs with less than 2 reads per million in the plasmid pool or a read count of zero at any time point are removed. | LFC across guides is scaled such that median of double non-targeting controls is set to 0 and median of positive control pairs (essential genes paired with non-targeting controls) is set to -1. | Expected DMF is modelled as the sum of SMF of each gene in the pair. GI score is calculated as the residual of each observed DMF from the control regression line. We refer to it as the Parrish score as we use the implementation provided by Parrish et al, but note that this method is a combination of techniques introduced in (19) and (20). | The R code was provided by the authors upon request and was adapted for each dataset. |

These scoring methods are applied on five different CDKO screens: the Thompson screen, the Parrish screen, the Ito screen, the Dede screen and the CHyMErA screen (Table 1). As each dataset includes distinct sets of gene pairs and cell lines, our approach cannot be used to compare the performance of different screens. Rather it allows us to assess how well each scoring approach performs on each individual screen.

The Gemini score is the only method that accounts for sample dependent as well as sample independent effects and therefore accounts for inherent variability and inter-dependencies in combinatorial screens. Sample independent effects account for various sources of variation across screens, such as differences in CRISPR guide activity, promoter strength, and batch effects. However, for studies with multiple cell-lines but without common early time-point read counts from plasmid pools (7, 9, 21), it is not possible to account for sample independent effects, as each cell line data has to be scored individually.

The key difference between Orthrus and other methods is that Orthrus takes orientation effects into account. This is specifically important for combinatorial CRISPR screens that use two different nucleases to target two genes and may have varying efficacies of guides at each position. An example of such screens is the CHyMErA screen (21), published by the same group and discussed in the next section, which uses Cas9 and Cas12a guides to target gene pairs. Even when a single nuclease is used at both positions, guides can have differential effectiveness depending on their position due to variation in promoter efficiency. Orthrus thus accounts for orientation, considering whether a gene is targeted by guide A at position A or by guide B at position B. It performs significance testing on expected and observed sets with matching orientations. However, this requires a reasonable number of guides in each orientation to gain statistical power. Orthrus can be configured to ignore orientation.

All studies, except the Parrish study and the Thompson study, target gene pairs in both orientations (A-B and B-A). The Parrish and the Thompson study target gene pairs in a single orientation. This breaks the Orthrus code which needs a gene-pair to be in both orientations. To address this, we implemented a workaround: for each gene pair, we duplicated the read counts (e.g., if there were initially 16 rows for each gene pair, there are now 32 rows) and swapped the positions of gene A with gene B, and guide1 with guide2. While this should not affect the final GI score, it may influence the statistical significance, which is not a primary concern in our analysis.

We also observed that the CAVI algorithm used in the Gemini score did not converge for either cell line analysed in the Parrish study. According to the Gemini user guide, the default initialization parameters must be adjusted to achieve convergence in such cases. However, at the time of writing this manuscript, the user guide does not provide further guidance on addressing this issue.

It is important to note that most methods outlined below use distinct filters for guides based on read counts or single gene essentiality. For our primary analysis, we omitted all such default steps to ensure consistent preprocessing across all methods and datasets. It is also worth noting that only Gemini and Orthrus are available in the form of R-packages, with comprehensive user guide documentation. This makes them easier to run compared to other methods. For all other methods, the code provided by the authors had to be adapted for each dataset.

### Benchmark Datasets

To evaluate each scoring method on the CDKO datasets, a benchmark set of pairs labelled as SL or non-SL is necessary. Since no established gold-standard exists for SL, we use two compiled lists of SL interactions derived from single-gene knockout CRISPR screens as benchmarks: De Kegel hits (17) and Köferle Hits (18). Both lists are derived from analysis of single-gene CRISPR screens across a panel of hundreds of cell lines and therefore should be robust across various cancer types and genetic backgrounds.

De Kegel Hits. This list contains SL annotations for 3,634 paralog pairs, with roughly 3.5% labelled as SL and the remainder not-SL. SL calls were made in (17) by analysing CRISPR screens in 768 cancer cell lines – a linear model was used to assess the association between loss of one gene and sensitivity to the inhibition of its paralog. Loss of a gene was based on three criteria: homozygous deletion, loss-of-function mutation or loss of mRNA expression. Due to a variety of restrictions on the paralog pairs tested (e.g. minimum sequence identity threshold, broad expression, gene loss in at least 10 cell lines) this dataset only annotates around 10% of the 36.6k protein coding paralog pairs from Ensembl used in De Kegel et al. (17). Each study in our analysis has a different proportion of overlapping gene pairs with this dataset as shown in Figure 2.

**Figure 2.**
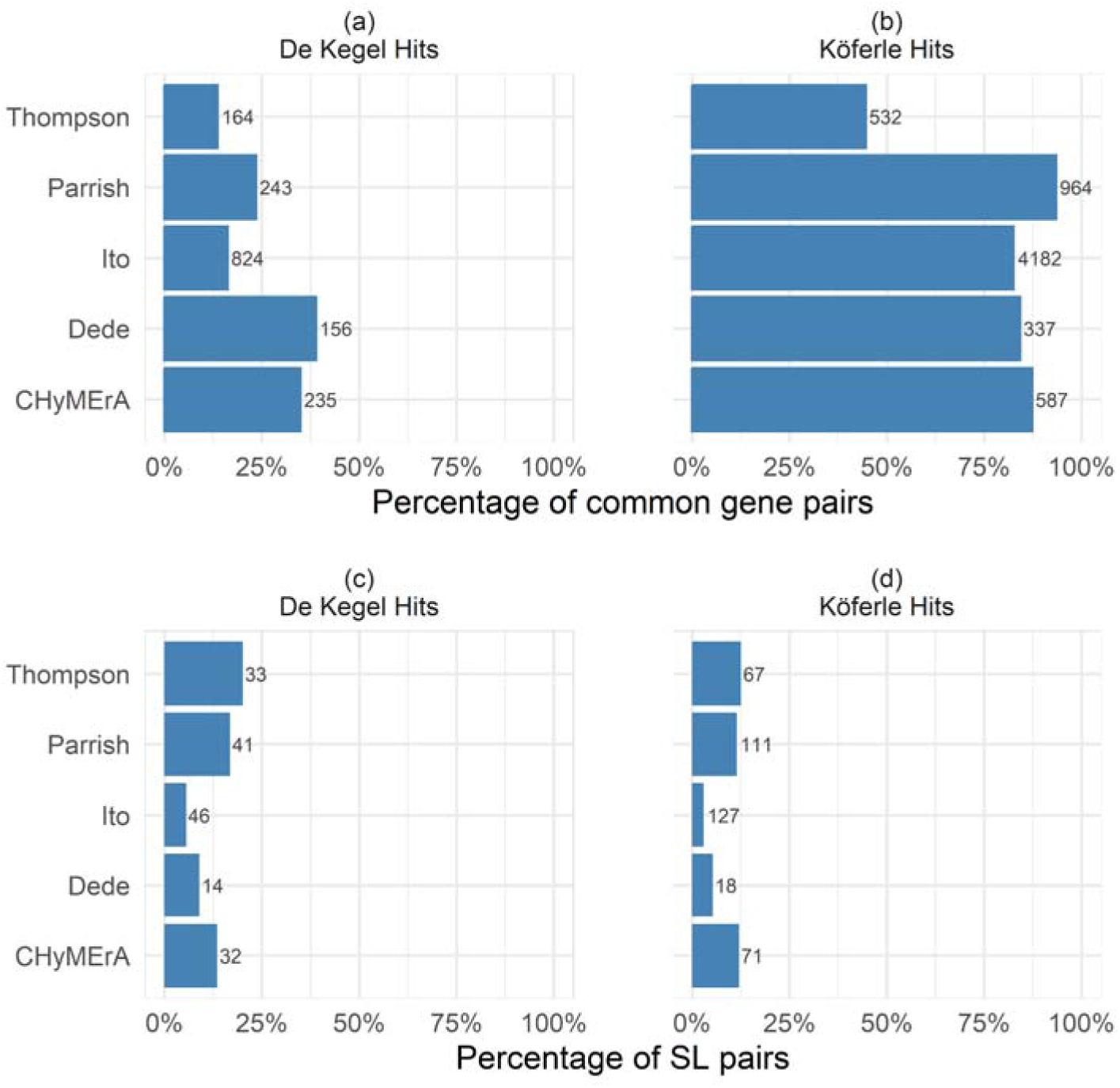
Analysis of the gene pairs in CRISPR DKO studies and benchmark datasets. (Top) Proportion of gene pairs in each study common with benchmark data (a) De Kegel Hits (b) Köferle Hits. Actual number of hits (rather than percentage) is indicated to the right of each bar. (Bottom) Percentage of gene pairs of each study that are identified as synthetic lethal in the benchmark datasets (c) De Kegel Hits (d) Köferle Hits.

Köferle Hits. This list contains annotations for 48,095 pairs, with 0.92% labelled as SL and the remainder not-SL. This list was generated using the same resources as the De Kegel et al. (17), however the associations were calculated by correlating the CRISPR dependency scores of one paralog and the mRNA expression of another. A ‘hit’ was identified when the Spearman’s correlation coefficient for a gene pair deviates by more than three standard deviations from the mean of all gene-pairs and its corresponding p-value was less than 0.05.

For our purposes, we define SL pairs as the subset of hits where the Spearman correlation is greater than 0 (indicating decreased expression of one results in increased dependency of the other). Non-SL pairs are those where the Spearman’s correlation coefficient is less than 0 and the pair is not labelled as a hit. For our study, we filter the pairs where the depletion/dependency score is taken from DepMap ‘AVANA’ screens (Meyers et al. 2017), as this dataset contains the largest proportion (46%) of the candidate biomarkers and is the most comprehensive in terms of the number of cell lines included. Compared to De Kegel Hits benchmark, this dataset has a higher percentage of overlapping gene pairs for each study as shown in Figure 2b.

Figure 2c and Figure 2d show the percentage of positive SL pairs out of total for both benchmark datasets. It is evident that all datasets are highly imbalanced, particularly the Ito study, where the share of positive SL pairs are significantly lower compared to others.

### Evaluation Metrics

To evaluate and compare the performance of each scoring method, we report two threshold-free measures: AUROC and AUPR. These metrics are computed using the trapezoidal rule with the Python scikit-learn package (version 1.5.2).

An ROC curve shows the trade-off between the false positive rate (FPR) and true positive rate (TPR) of a binary classifier (22). AUROC of 0.5 indicates a random classifier. For imbalanced datasets, like we have, the PR curve is often more informative than the ROC (23) as it shows a trade-off between precision and recall (TPR) across varying threshold settings. The baseline performance in a PR curve is represented by a horizontal line equal to the ratio of positive samples to total samples.

### Default filtering settings of GI scoring methods

As noted in Table 2, each method returns a different number of scored pairs due to varying default filtering settings. To ensure a fair comparison, we standardized the methods by removing these pre-processing steps, forcing them to score all pairs regardless of low read counts or whether one of the genes might be essential. The Gemini-Sensitive essentiality filter removed 10-20% of gene pairs in all screens except the Dede screen and Orthrus default filter removed almost 50% of the pairs in the CHyMErA screen (Figure S1). Therefore, we also include an analysis of how the filtering steps of the Gemini-Sensitive score and the Orthrus score impact the overall results.

## RESULTS

### All scoring methods are correlated

We first applied each scoring system to the results of each screen in each dataset, resulting in a total of 105 scored datasets encompassing 21 individual cell line screens. This allowed us to assess the agreement between scores generated by different approaches, presented in Figure 3, aggregated over all screens, with the comparisons across individual screens presented in Figure S2. As might be expected, the scores generated by all scoring approaches are at least moderately correlated (minimum Spearman correlation of 0.42) (Figure 3a). Two clear groupings emerge from this analysis – one group containing the two Gemini variants (sensitive and strong) and a second group containing the three other scores (Orthrus, Parrish and zdLFC). Gemini is conceptually different from the other three scores in that it uses a Bayesian approach to estimate single and combination effects while accounting for guide specific variation in efficiency and cell line specific effects. Genetic interactions are then scored using these inferred combination and single gene effects, with the difference between the strong and sensitive scores being in how they are subsequently processed. In contrast the other three methods primarily model the expected DMF as a linear function of the observed SMF and compare this expected value with the observed DMF.

**Figure 3.**
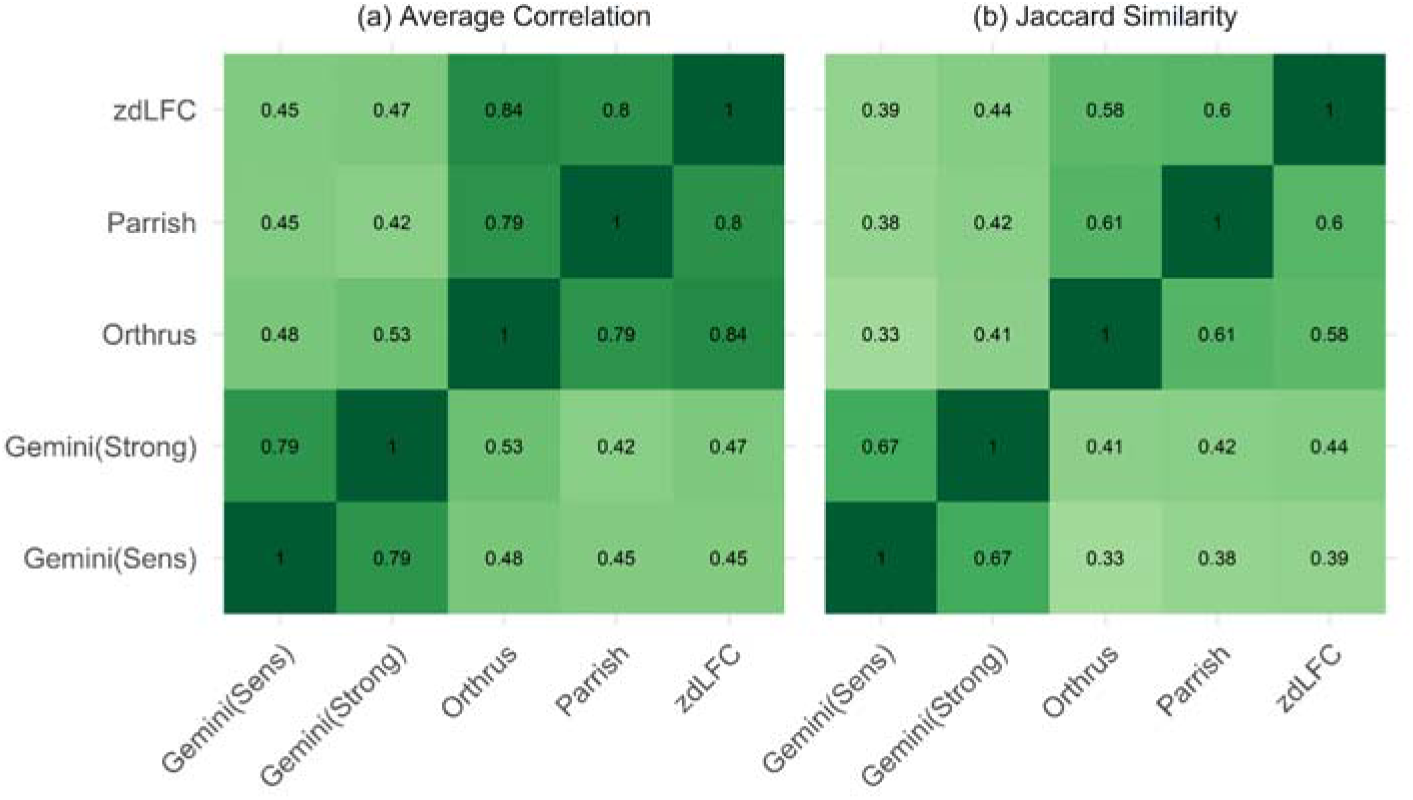
Agreement between scores calculated by different methods. (a) Average correlation across methods for all studies, (b) Average Jaccard similarity of top 5% scored gene pairs across methods. Darkest shade of green shows an average correlation of 1 or average Jaccard index of 1.

As only a minority of gene pairs are expected to display genetic interactions, and hence most scores may be close to zero, we also analysed the agreement between the top 5% of gene pairs identified by each scoring system (i.e. the pairs with the most negative genetic interaction effect, indicative of synthetic lethality). To quantify the agreement between scoring systems we used the Jaccard index – the size of the intersection between the hits identified by both systems divided by the size of the union of the hits, with a value of 0 indicating no overlap and 1 indicating complete overlap (Figure 3b). Consistent with our correlation analysis, the scoring systems fell into two distinct groups with Gemini-Sensitive and Gemini-Strong scores in one group with high Jaccard scores between them and Orthrus, zdLFC and Parrish in the other. While there is variation in the average Jaccard across different studies (Figure S2b) these groupings are generally consistent.

### The Parrish score consistently yields higher AUROC, and the Gemini-Sensitive score yields higher AUPR compared to others using De Kegel Hits as the benchmark

We next analysed the performance of each scoring system across each study using the benchmark datasets. For our analysis, we computed AUROC and AUPR metrics by combining all cell lines within each study, and also by analysing each cell line individually within each study. We focus primarily on performance aggregated by study rather than by cell line, to avoid a single study (11) with a large number of cell lines dominating the results. We then assessed the performance of each score-study combination, as well as each score-cell line combination using the two benchmarks.

Evident from Figure 4a, the Parrish score achieves the highest AUROC in three of the studies: the Thompson study, the Parrish study and the Dede study. Gemini-Sensitive score yields the highest AUROC for the Ito study and the Orthrus score yields the highest AUROC for the CHyMErA study. We note that the Gemini score was developed by authors of the Ito study, Orthrus was developed by the authors of the CHyMErA study, zdLFC was developed by the authors of the Dede study, while Parrish et al. (9) adapted previous scoring systems to apply to their own screens. In general therefore, the highest AUROC is observed when the method developed by a group of authors is paired with the screen performed by those authors.

**Figure 4.**
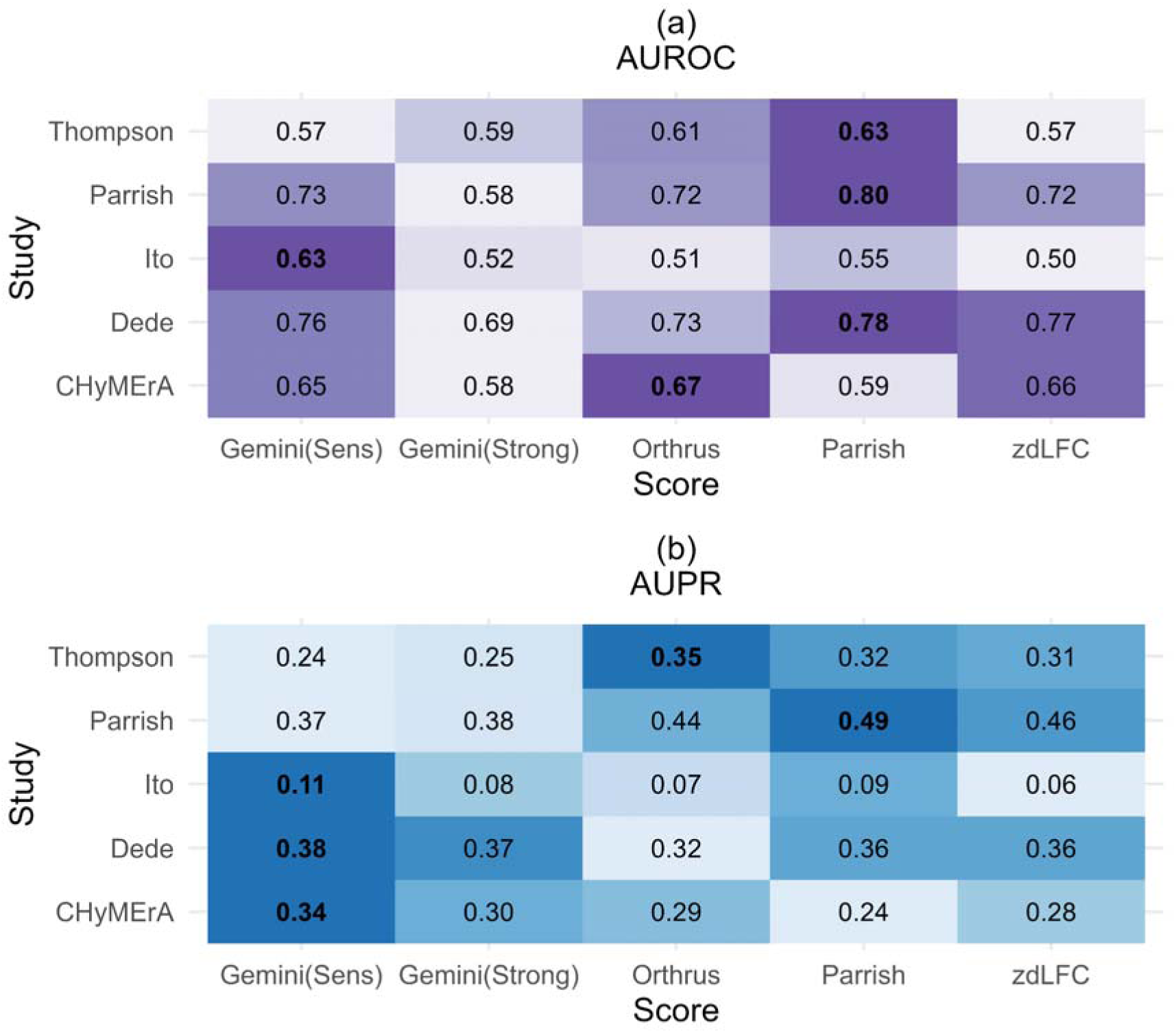
Evaluation of scoring systems using the De Kegel Hits as a benchmark. (a) AUPR and (b) AUROC of each study with all cell lines combined. The colour represents the normalized AUROC/AUPR for each study, with the darkest shade of purple/blue indicating the highest AUROC/AUPR within the corresponding study.

As shown in Figure 4b, the Parrish score achieves the highest AUPR in one out of five datasets (Parrish), whereas Gemini-Sensitive leads in three (Ito, Dede and CHyMErA).

When analysing individual cell lines, the Gemini-Sensitive score yields the highest AUROC for 12 out of 21 cell lines (primarily those from the Ito study), followed by the Parrish score that yields the highest AUROC for 6 out of 21 cell lines (primarily those from Parrish, Dede and Thompson) (Figure S3). The Gemini-Sensitive score yields highest AUPR for 11 (primarily Ito), the Parrish score yields highest AUPR for 5 cell lines (including both screened by Parrish) and the Orthrus score yields the highest AUPR for 4 cell lines (Figure S3).

### The Gemini-Sensitive score consistently yields higher AUROC and AUPR compared to others using Köferle Hits as the benchmark

Figure 5a shows the Gemini-Sensitive scores achieving the highest AUROC and AUPR in three studies (Ito, Dede and CHyMErA), the Parrish score achieves the highest AUROC and AUPR in two studies (Thompson and Parrish) (Figure 5b). These results are mirrored when analysing individual cell lines, where the Gemini-Sensitive score has generally the highest AUPR and AUROC scores in cell lines from the Ito, Dede and CHyMErA screens, and the Parrish score has the highest AUPR and AUROC scores for those from Parrish and Thompson screens (Figure S4).

**Figure 5.**
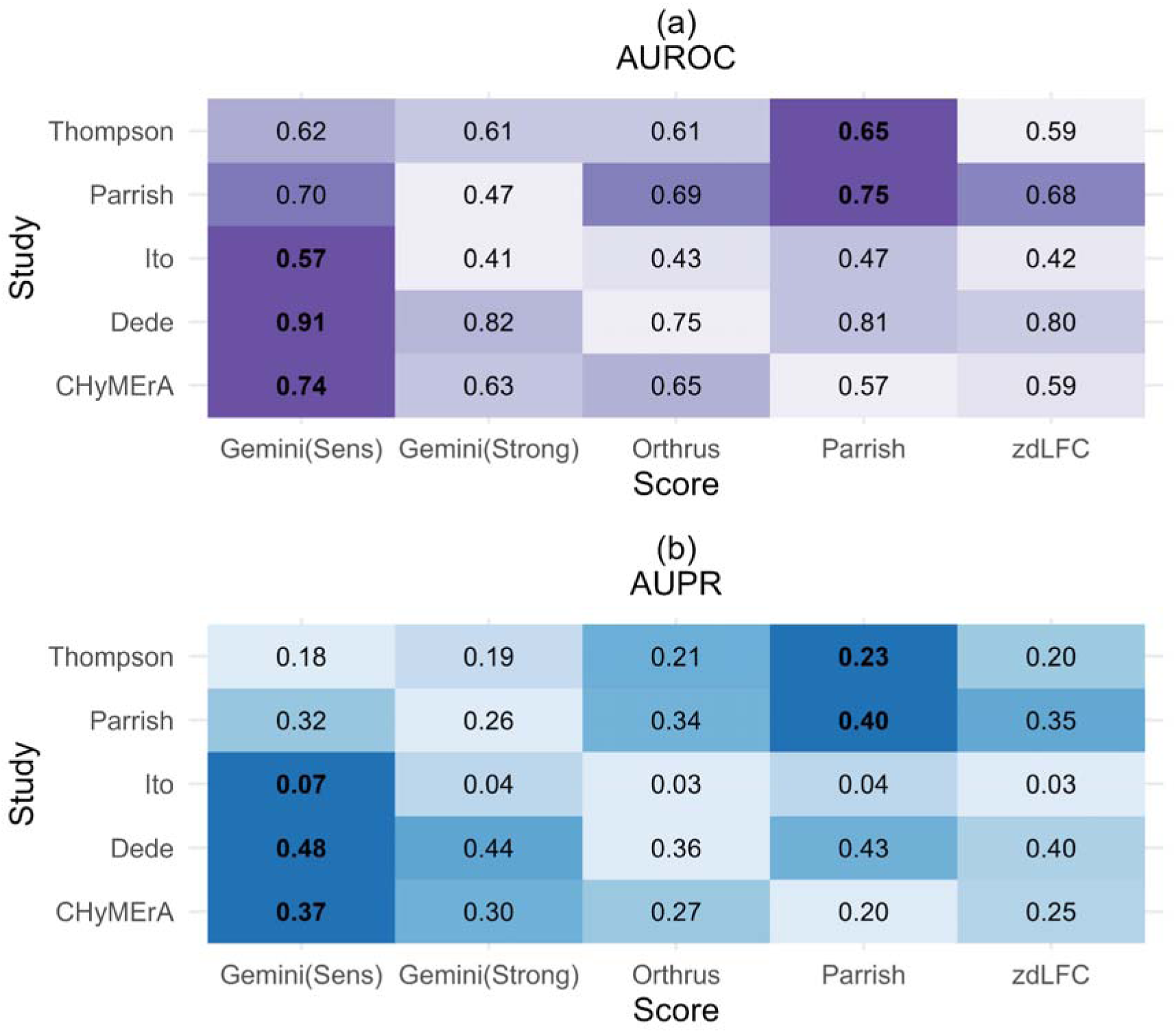
Evaluation of scoring systems using the Köferle Hits as a benchmark. (a) AUPR and (b) AUROC of each study with all cell lines combined. The colour represents the normalized AUROC/AUPR for each study, with the darkest shade of purple/blue indicating the highest AUROC/AUPR within the corresponding study.

The Parrish score performs best on the Parrish screen and the Thompson screen using both benchmarks with respect to AUROC (Figure 4-5). In case of AUPR, the Parrish Score performs best for the Parrish screen when De Kegel hits are used as benchmarks and performs best both for the Thompson screen and the Parrish screen with Köferle hits are used as benchmarks. Both studies have an asymmetric guide layout, meaning that the combination of genes A and B is knocked out in only one orientation (A-B), with no (B-A) orientation, suggesting that the Parrish score may be the best scoring system in this setting.

### The default essential gene filter in Gemini-Sensitive score improves results

The default filtering step in the Gemini-Sensitive score drops a significant number of gene pairs (between 0.4% to 20%) across the studies (Figure S1). We assessed the impact of this filtering on AUROC and AUPR for each study. AUROC and AUPR values of the filtered datasets are generally higher compared to the unfiltered datasets (Figure 6a and 6b), when evaluated against both benchmarks. This indicates that the single-gene essentiality filter helps to reduce false positives and hence improves AUPR and AUROC. However, the overall improvement is modest, with a maximum increase of 5% for AUROC and 3% for AUPR.

**Figure 6.**
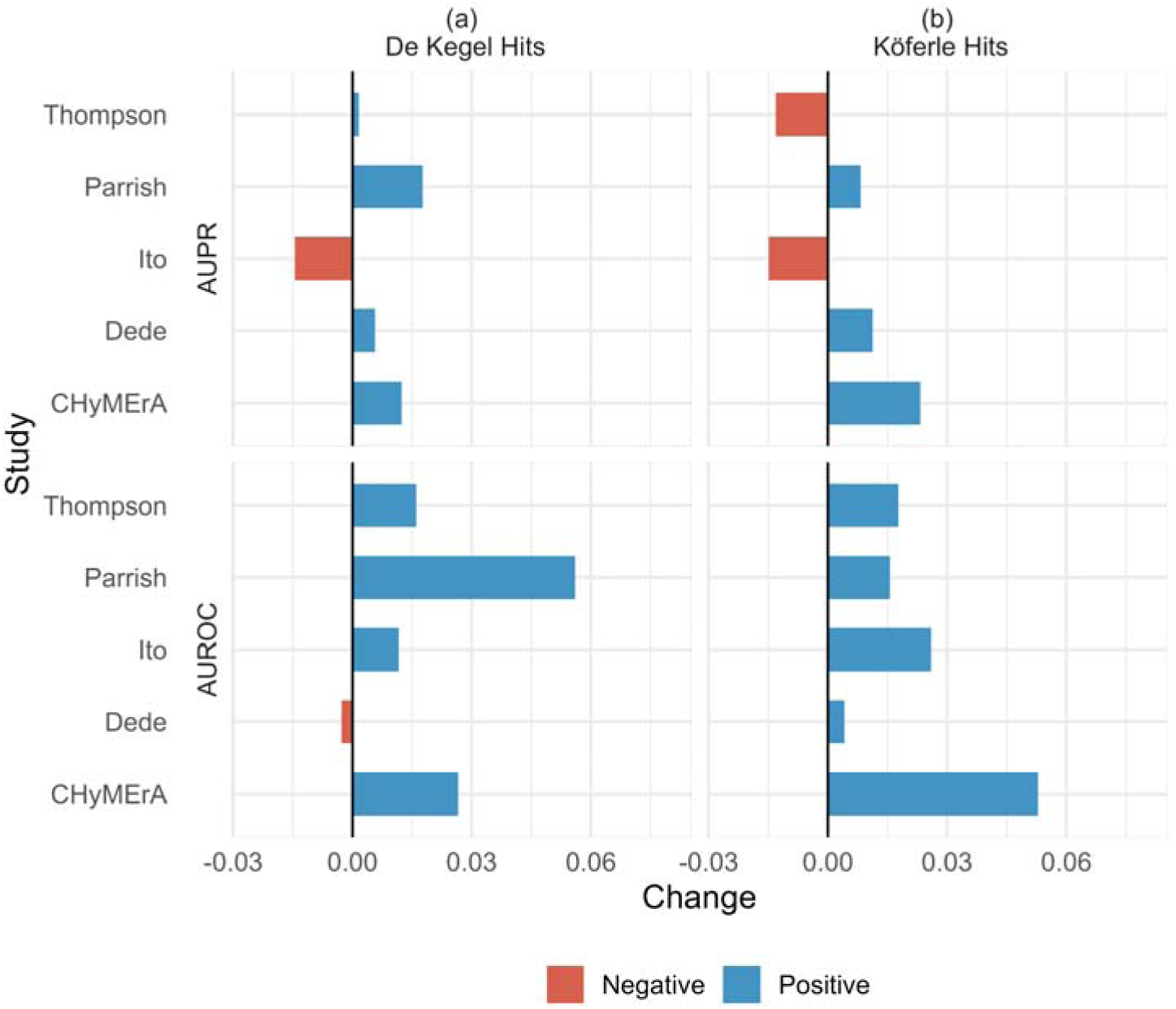
Gemini filtering step generally improves performance. Panels (a) and (b) show the changes in AUPR and AUROC for each study when the essential gene filter of Gemini-Sensitive scoring method is applied versus not applied, using (a) De Kegel Hits and (b) Köferle Hits as benchmarks. A positive value indicates that the filtering step improves the results.

### The default read count filter of Orthrus discards usable information

The Orthrus default filtering steps result in almost 50% genes from the CHyMErA dataset being excluded from analysis (Figure S1). We compared the AUROC and AUPR of gene pairs that were filtered out (n = 236) with those that were not (n = 234) (Figure 7a and Figure 7b). Interestingly, while the AUROC of both sets was similar, the AUPR of the filtered-out gene pairs was significantly higher than that of the retained pairs. This suggests that removing gene pairs based on a small number of guides remaining post-filtering may discard valuable information, as these filtered-out pairs still perform well on both benchmarks. Similar observations were made when Köferle Hits was used as the benchmark (Figure 7c and 7d).

**Figure 7.**
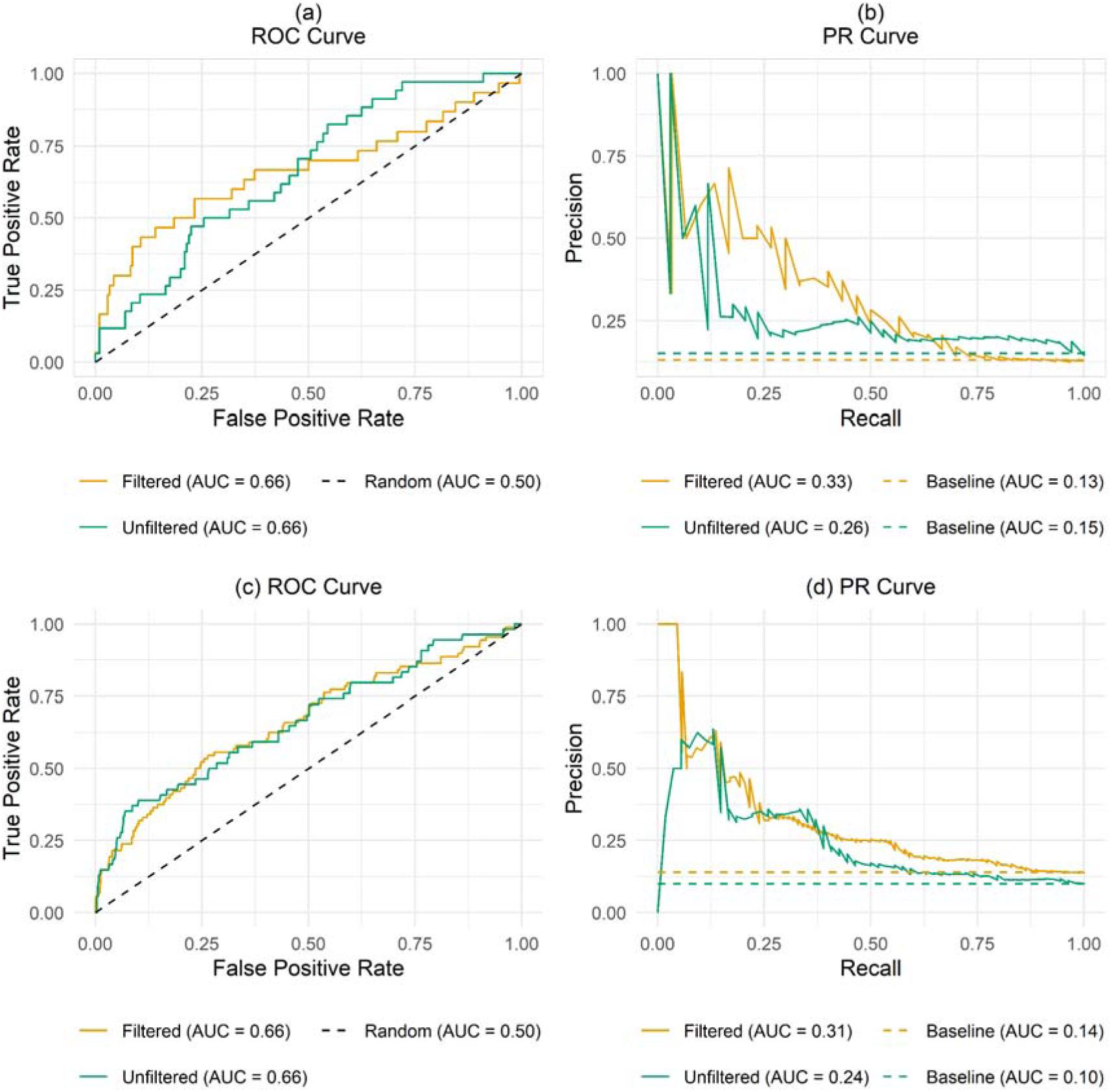
Impact of Orthrus default filtering steps. (a) ROC curve and (b) PR Curve of Orthrus score on CHyMErA pairs that are filtered out (n = 236) in default filtering step versus CHyMErA pairs that are not filtered (n = 234) out using De Kegel Hits as benchmark. (c) ROC curve and (d) PR Curve of Orthrus score on CHyMErA pairs that are filtered out (n = 636) in default filtering step versus CHyMErA pairs that are not filtered (n = 538) out using Köferle Hits as benchmark.

## DISCUSSION

Identifying synthetic lethality (SL) is invaluable in cancer therapy research, however determining the most suitable statistical method to accurately identify SL from a CRISPR screen remains highly challenging.

Here, we have performed the first systematic evaluation of the existing GI scoring methods for CDKO screens. We have used 5 different scoring methods on 5 different CRISPR DKO studies and evaluated the performance of each scoring method on each study using two benchmarks. We found that the results from all methods exhibit moderate to strong correlation. In particular, Gemini-sensitive is highly correlated with Gemini-strong, and both Gemini variants have comparatively moderate correlation with rest of the methods. The other three methods—zdLFC, Parrish, and Orthrus—model the expected LFC as a linear combination of observed individual gene knockout effects. In contrast, Gemini employs a variational Bayesian approach to model the observed LFC as a function of both sample-independent and sample-dependent individual effects of each guide in a pair and the effect of two guides in combination. The Gemini-strong variant compares the combination effect of a gene pair to the most pronounced individual effect (flags an interaction when the combination effect exceeds the effect of either gene), while Gemini-sensitive compares the total effect (the sum of the individual gene effects and the combination effect) to the most lethal individual gene effect.

In most scenarios, either the Parrish score or the Gemini sensitive score yields the highest AUROC and AUPR (Figure 4 and Figure 5). The Parrish score gives superior results on datasets which use an asymmetric guide orientation and should therefore be used on such datasets. However, if we consider user-friendliness, installation procedures, code availability, reproducibility, and documentation quality as secondary measures (24), Gemini-Sensitive is more user friendly compared to the Parrish score as it comes as an easy-to-use R package with comprehensive documentation. One current drawback of using the Gemini score is that no instructions are available to adjust the model initialisation parameters for the models that do not converge with default parameters.

We also conduct a secondary analysis on the impact of default filtering steps of the Gemini-Sensitive and the Orthrus methods. We found that in the case of Gemini-Sensitive score, filtering essential gene pairs as recommended by the developers prior to scoring results in an increase of AUPR and AUROC in most studies (Figure 6). We also found that the default filtering step of the Orthrus scoring system discards usable information, i.e. AUPR of unfiltered scored pairs is higher compared to AUPR of filtered scored pairs (Figure 7).

Here we have evaluated combinatorial CRISPR screens that assess synthetic lethality between carefully selected gene pairs – paralog pairs. In this scenario, each gene may only be assessed for genetic interaction with one other gene (its paralog) and hence the estimation of single gene effects is heavily dependent on control constructs. In contrast to this approach, others have made use of matrix screens, where sets of up to hundreds of genes are screened in an all-against-all fashion Horlbeck et al. (25). Because every individual gene in these studies is tested in hundreds of combinations, it may be possible to gain an improved estimate of the size and variance of single gene effects from all double perturbations rather than relying on estimates from a small number of control constructs, as has been done in yeast (26). Therefore, it may also be worth evaluating different scoring systems in this setting, but the challenge will be the identification of a suitable benchmark.

## DATA AVAILABILITY

All analysis and code for reproducing the results of the study are publicly available at https://github.com/cancergenetics/Benchmarking-GI-Scores. The results of applying each scoring method to each screen are available at https://figshare.com/s/59ee190b1879fe3eb191.

## SUPPLEMENTARY DATA

Supplementary Data are available at NAR online (attached with the submission).

## Supporting information

Supplementary Images

## ACKNOWLEDGEMENTS

We would like to acknowledge Birkan Gökbağ, one of the authors of the SLKB database (16) and Alice H Berger, corresponding author of Parrish et al. (9), for assisting with code execution. We also acknowledge the Research IT HPC Service at University College Dublin for providing computational facilities and support that contributed to the research results reported in this paper.

