## Supplementary Images for "Benchmarking Genetic Interaction Scoring Methods for Identifying Synthetic Lethality from Combinatorial CRISPR Screens"

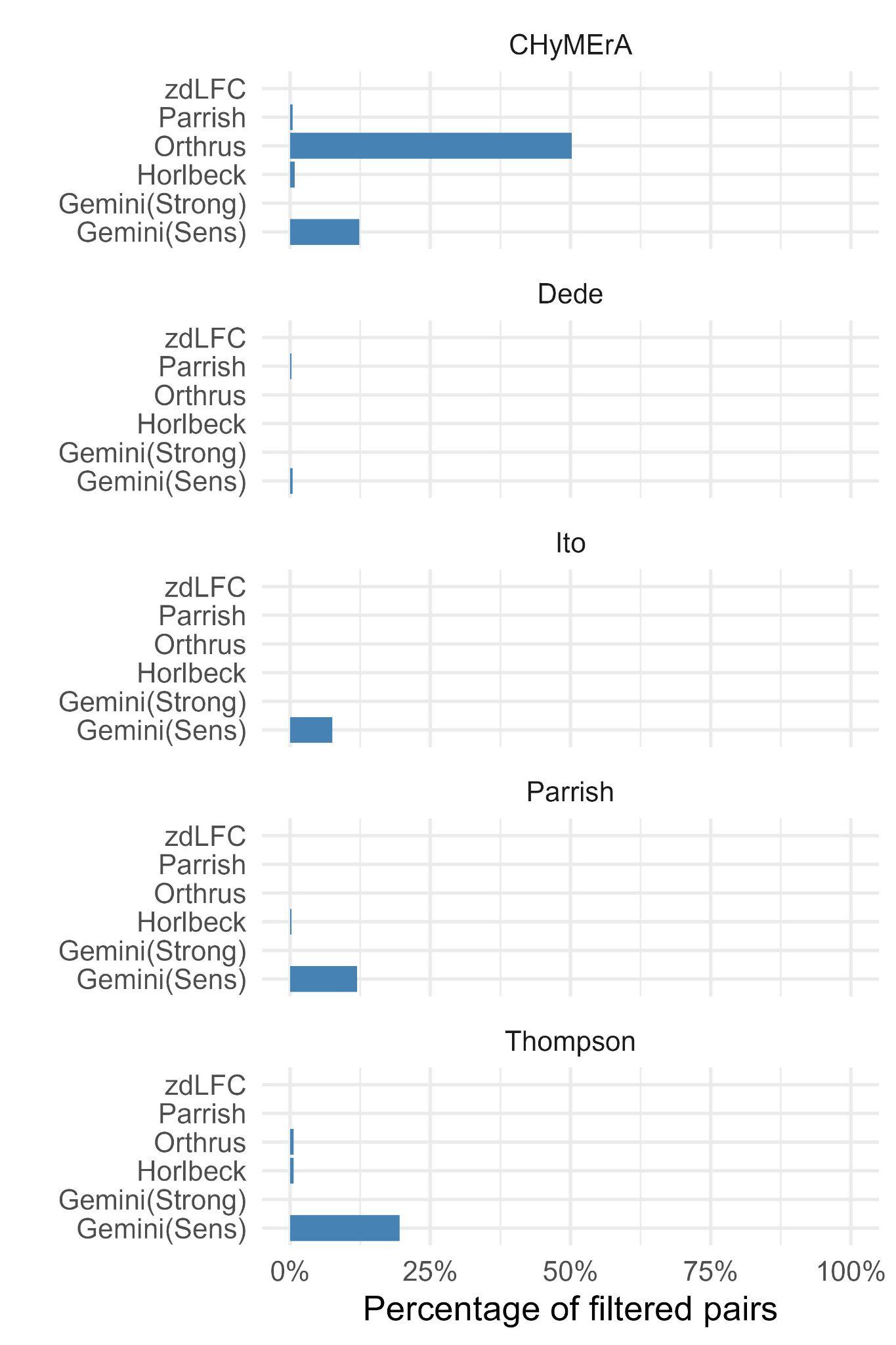

**Figure S1**. Percentage of gene pairs filtered by each method when default filters are applied.

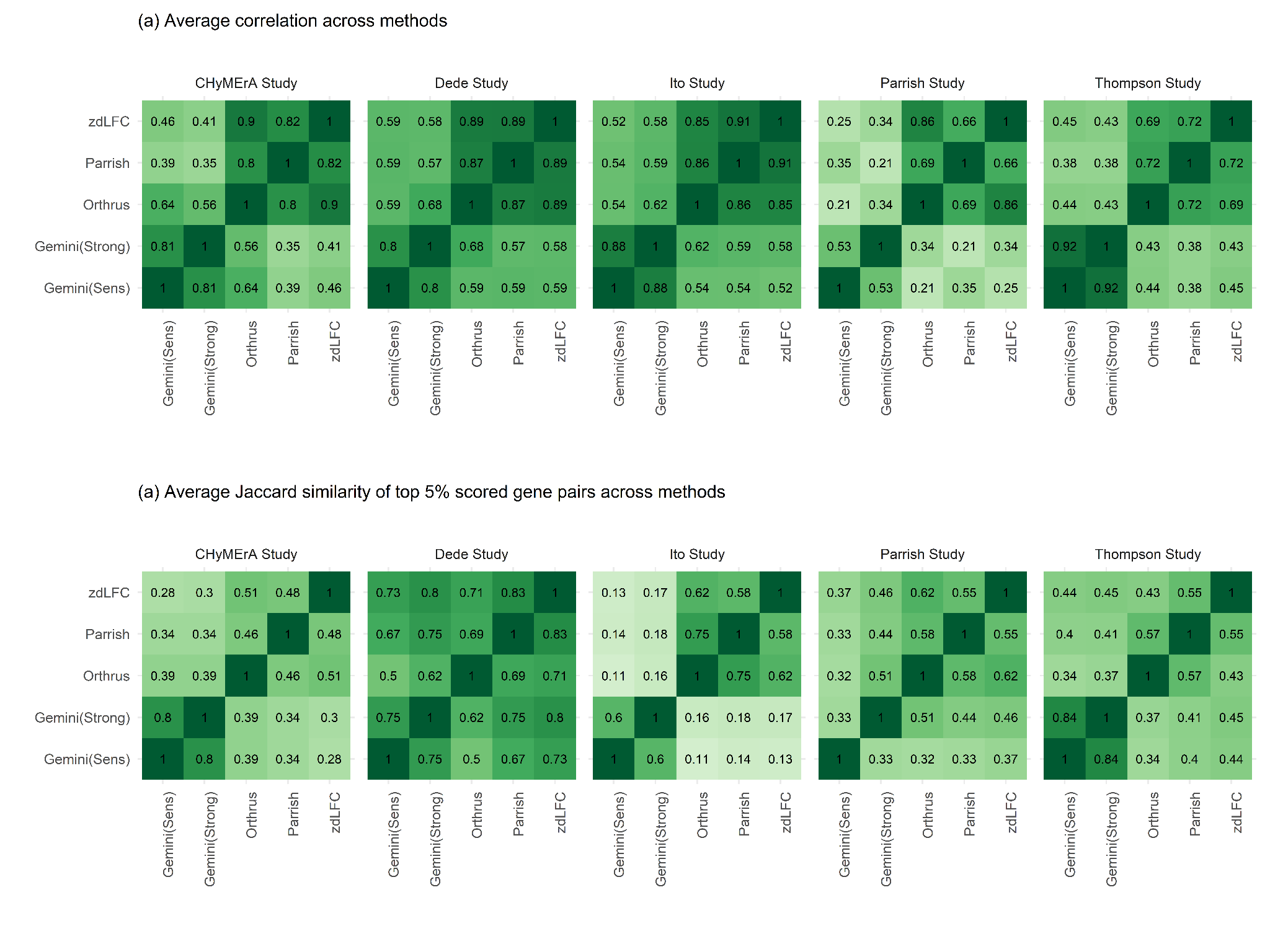

**Figure S2.** Comparison of Scoring Methods (a) Average correlation of scoring methods within each study; (b) Average Jaccard similarity of the top 5% scored gene pairs across scoring methods within each study.

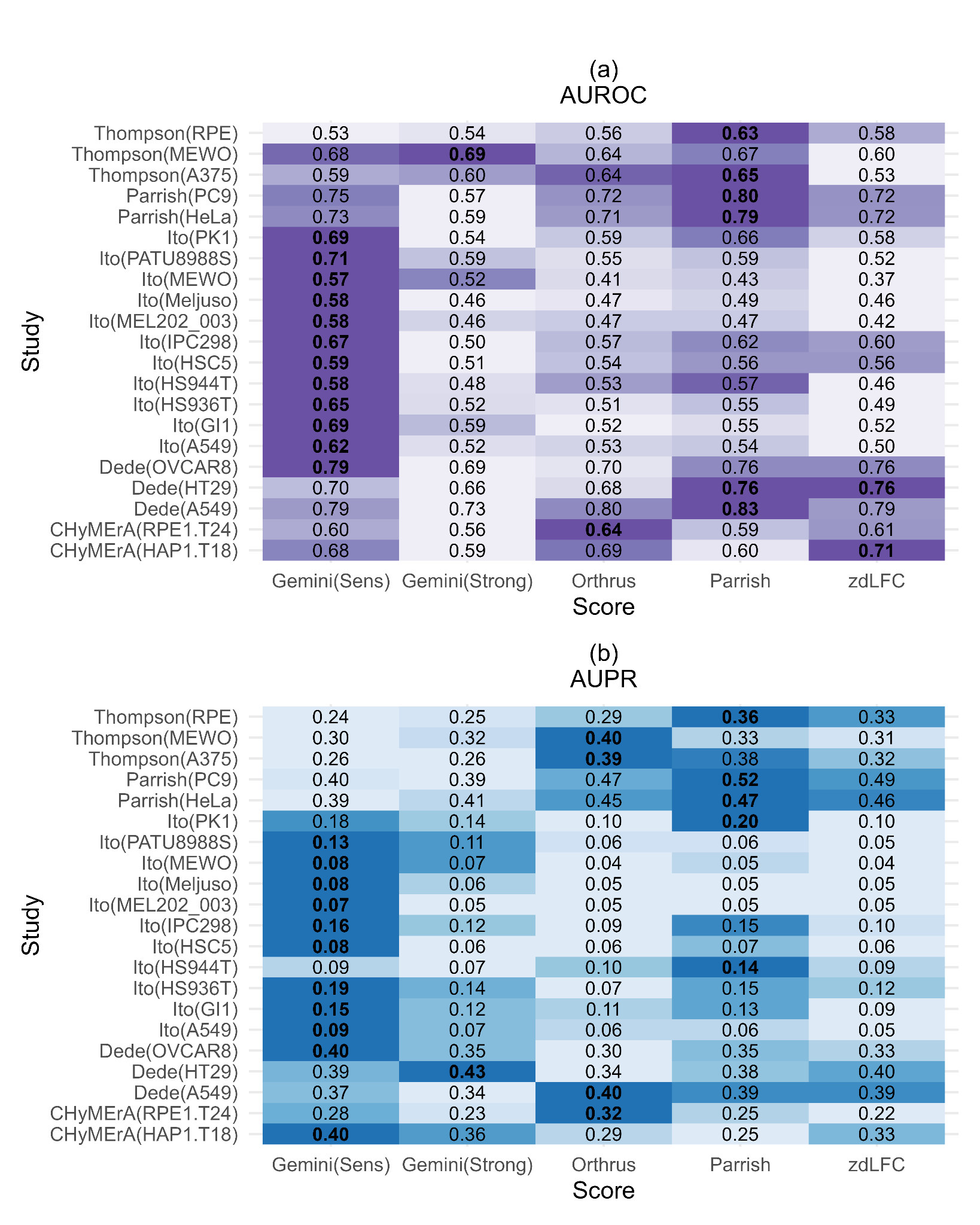

**Figure S3.** Evaluating scoring systems using the De Kegel Hits benchmark. (a) AUROC and (b) AUPR across individual cell lines of each study using De Kegel Hits as benchmark.

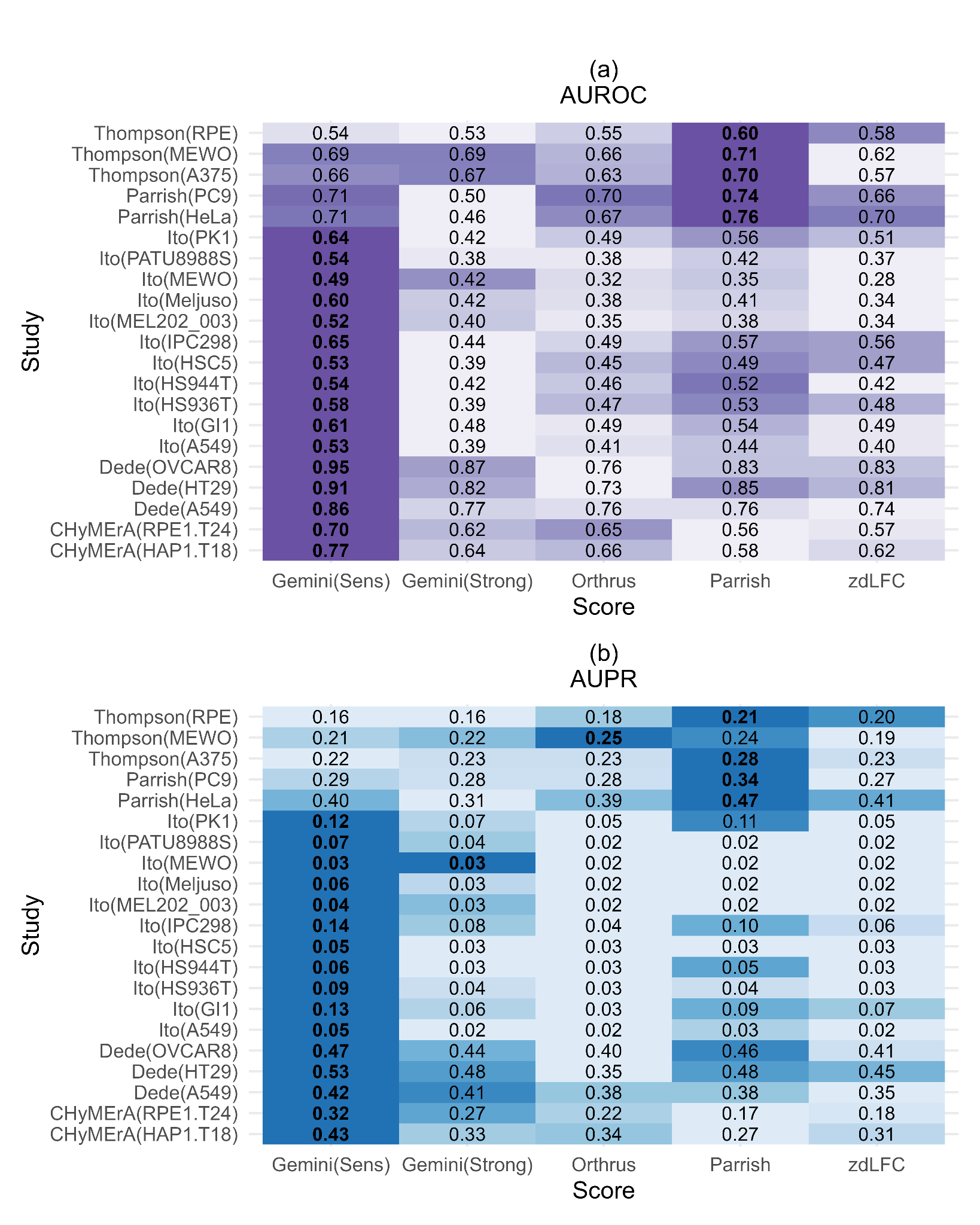

**Figure S4.** Evaluating scoring systems using the Köferle Hits benchmark. (a) AUROC and (b) AUPR across individual cell lines of each study using Köferle Hits as benchmark.
